# Residual DNA in enzymes may pose a challenge for DIY RNA sequencing protocols: how to deal

**DOI:** 10.1101/2022.02.02.478843

**Authors:** Andrey Krivoy, Yegor Botsmanov, Valery Cheranev, Anna Shmitko, Vera Belova, Viktoriya Moskalenko, Robert Afasizhev, Anastasia Shut, Dmitriy Korostin, Denis Rebrikov

## Abstract

High-throughput RNA sequencing (RNA-seq) is widely employed for gene expression profiling. This report on do-it-yourself RNA-Seq library preparation pitfalls, highlighting their correct identification and successful overcoming, may be helpful to research groups commencing a DIY RNA-Seq workflow and facing similar problems.

## Introduction

High-throughput RNA sequencing (RNA-seq) is widely employed for gene expression profiling. A typical RNAseq workflow is straightforward as it relies on optimized commercial kits. The major drawback of ready-to-use kits for RNA-seq library preparation is their high costs per sample. For RNAseq demanded on a routine basis, do-it-yourself (DIY) library preparation protocols may represent an advisable option. As all steps of the conventional RNA-seq library preparation workflows are well understood and optimizable, these workflows can be successfully reproduced with separately purchased or home-made reagents without any proprietary concerns. Apart from a significant reduction in the costs per sample, the DIY protocols can be readily adjusted for biological samples with specific properties (low RNA amounts, suboptimal degree of RNA fragmentation, admixtures of RNA from different species, etc.). In addition, the use of DIY workflows is beneficial for staff qualification and affords better control over the library preparation routine. Here we describe our experience with a newly designed DIY RNA-seq library preparation protocol. Starting with low amounts of RNA, we encountered a bias related to contamination of recombinant enzymes with residual host-cell DNA. This report on DIY RNA-Seq library preparation pitfalls, highlighting their correct identification and successful overcoming, may be helpful to research groups commencing a DIY RNA-Seq workflow and facing similar problems.

## Results

### Detection of DNA contamination traced to enzymes

A newly designed DIY RNA-seq protocol (see Methods) was tested on serial dilutions of RNA extracted from HEK293 human cells. The data was mapped using STAR aligner [1] under standard parameters; multi-mapped reads were recorded for every mapped region (match) with a top MAPQ score and the counts were divided equally. Visual inspection of the NGS data mapped to human genome confirmed the domination of spliced mRNA sequences expected for the poly(dT) oligo-primed reverse transcription (Figure 1). Main metrics for the sequencing data composition are presented in Table 1. As can be seen, the ratios between exome/introns/rRNA/lncRNA and intergenic regions did not react dramatically to the decreasing amounts of input RNA, while the percentage of reads not mapped to human genome significantly increased with the decrease of RNA input.

**Figure 1.**
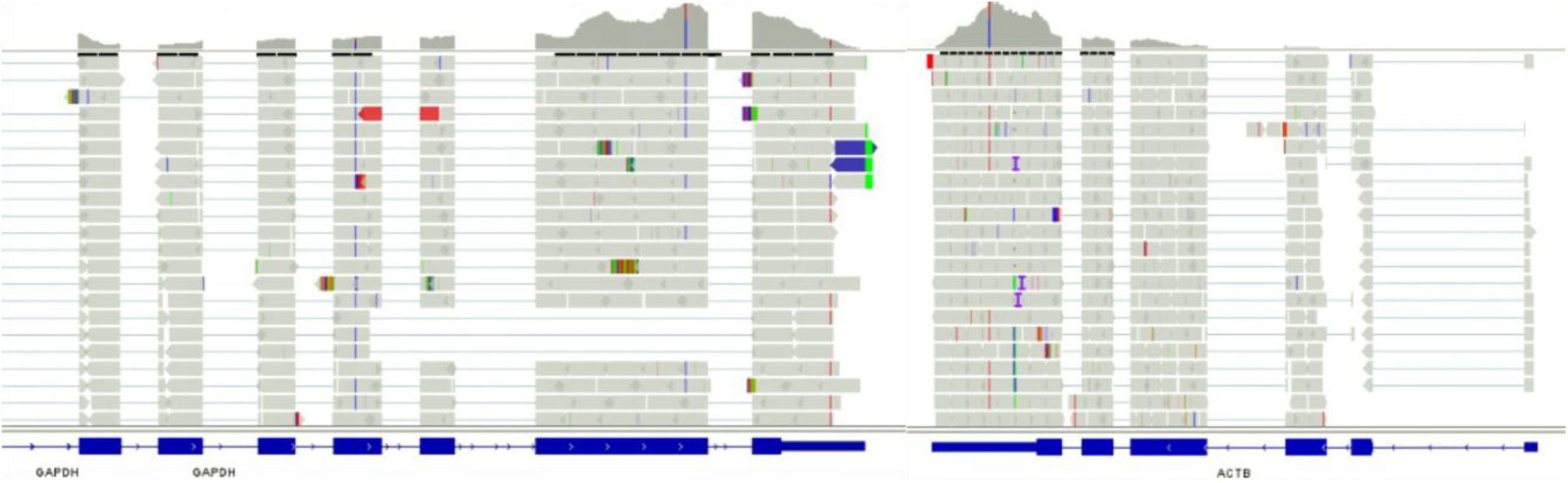
An Integrative Genomics Viewer (IGV) [2] chart confirming the domination of spliced mRNA sequences among gene-covering RNAseq data. Two housekeeping genes, *GAPDH* and *ACTB*, were selected for a representative pattern. The sequenced mRNA is mostly spliced, as polyA tailing, which provides the tag for the enrichment, occurs at an advanced stage of mRNA maturation [3].

**Table 1.**
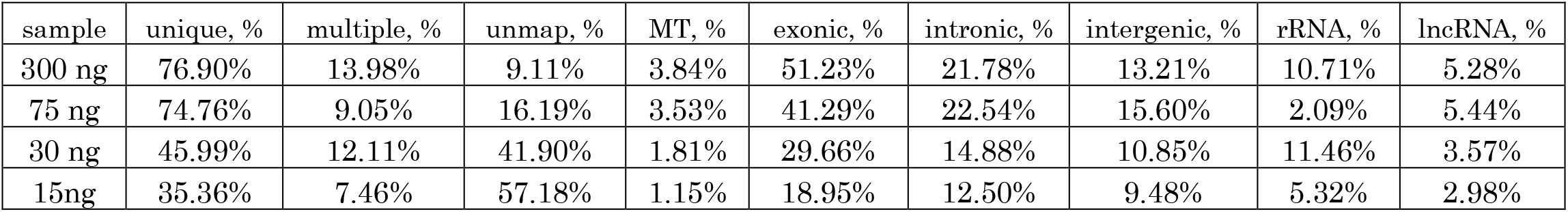
NGS metrics for RNA extracted from HEK293 cells. Uniquely mapped reads were identified by STAR aligner as having a unique location in human genome. Multiple-mapped reads were aligned to at least two different genomic locations each. Unmapped reads showed no significant similarity to human genome. MT index reflects the percentage of mitochondrial genome transcripts in the data. Exonic, intronic, intergenic, rRNA and lncRNA reads match corresponding regions in human genome; exonic regions include rRNA sequences. The percentage of unmapped reads shows a clear negative correlation with the amount of RNA at the start.

| sample | unique, % | multiple, % | unmap, % | MT, % | exonic, % | intronic, % | intergenic, % | rRNA, % | lncRNA, % |
| --- | --- | --- | --- | --- | --- | --- | --- | --- | --- |
| 300 ng | 76.90% | 13.98% | 9.11% | 3.84% | 51.23% | 21.78% | 13.21% | 10.71% | 5.28% |
| 75 ng | 74.76% | 9.05% | 16.19% | 3.53% | 41.29% | 22.54% | 15.60% | 2.09% | 5.44% |
| 30 ng | 45.99% | 12.11% | 41.90% | 1.81% | 29.66% | 14.88% | 10.85% | 11.46% | 3.57% |
| 15ng | 35.36% | 7.46% | 57.18% | 1.15% | 18.95% | 12.50% | 9.48% | 5.32% | 2.98% |

The standard parameters of STAR conventionally used for the analysis require 66 % of the reads to be matched to the reference genome to declare the species identity. The unmapped reads in sequencing data may have no relation to human transcriptome/genome and represent completely different organism(s), probably commensal or contaminant. We explored this situation using Kraken2 taxonomic sequence classifier [4] with a k-mer matcher applied throughout a phylogenomic database. As expected, a major fraction of reads identified with *Homo sapiens*. Surprisingly, a fraction approximately equal to the unmapped-by-STAR counterpart identified with *Escherichia coli* genomic sequences along with T4-like viral sequences (Table 2). To classify the issue as biological *vs*. technical, we addressed the no-template negative control (NTC) library preparation with the same DIY protocol. Surprisingly, at 16 PCR cycles we obtained libraries with detectable concentrations above the Qubit HS dsDNA sensitivity threshold and observable by agarose gel electrophoresis. Combined analysis by Kraken2 and BWA-MEM unambiguously identified *E. coli* and T4 genomic sequences accounting for more than 80 percent of the total data. This finding directly implicated a certain source of DNA contamination apparently related to the DIY RNA-seq library preparation workflow. We assumed that certain recombinant enzyme(s) used in the workflow, produced in bacterial host cells transformed with vectorized coding sequences for the protein production, might be contaminated.

**Table 2.** Mapped-to-species characteristics of RNAseq libraries made from RNA dilutions. Kraken2, STAR, and BWA-MEM output for the libraries prepared from RNA samples (Table 1) and a no-template negative control (no cDNA or gDNA input, NTC). Each value represents the number of reads identified with particular organism (taxon) divided by the total number of reads. The Kraken2 output is almost identical to the corresponding STAR and BWA-MEM values. The higher RNA concentrations are used, the lower is the degree of contamination with *E. coli* (Enterobacteriaceae) and Escherichia virus T4 (*Tequatrovirus*, Myoviridae) sequences.

| Sample | NTC | 15ng | 30ng | 75ng | 300ng |
| --- | --- | --- | --- | --- | --- |
| Kraken2 *Homo sapiens*, % | 0 | 45 | 60.1 | 85.3 | 92.8 |
| Kraken2 Enterobacteriaceae, % | 48.4 | 27.61 | 19.8 | 6.21 | 2.1 |
| Kraken2 Tequatrovirus, % | 31.8 | 11.06 | 7.47 | 2.87 | 1 |
| STAR mapped to human, % | N/A | 42.82 | 58.1 | 83.8 | 90.1 |
| BWA mapped to *E. coli*, % | 45.4 | 26.85 | 19.35 | 5.96 | 2.04 |
| BWA mapped to T4 genome, % | 31.5 | 11.6 | 7.91 | 3.01 | 1.06 |

### Identification of contaminated enzyme

To identify the exact source of DNA contamination in the DIY workflow, we mapped sequencing data for the negative control and “15 ng” (Table 1) libraries to *E. coli* and T4 phage reference genomes (Figure 2, Table 2) using BWA-MEM with default parameters [5].

**Figure 2.**
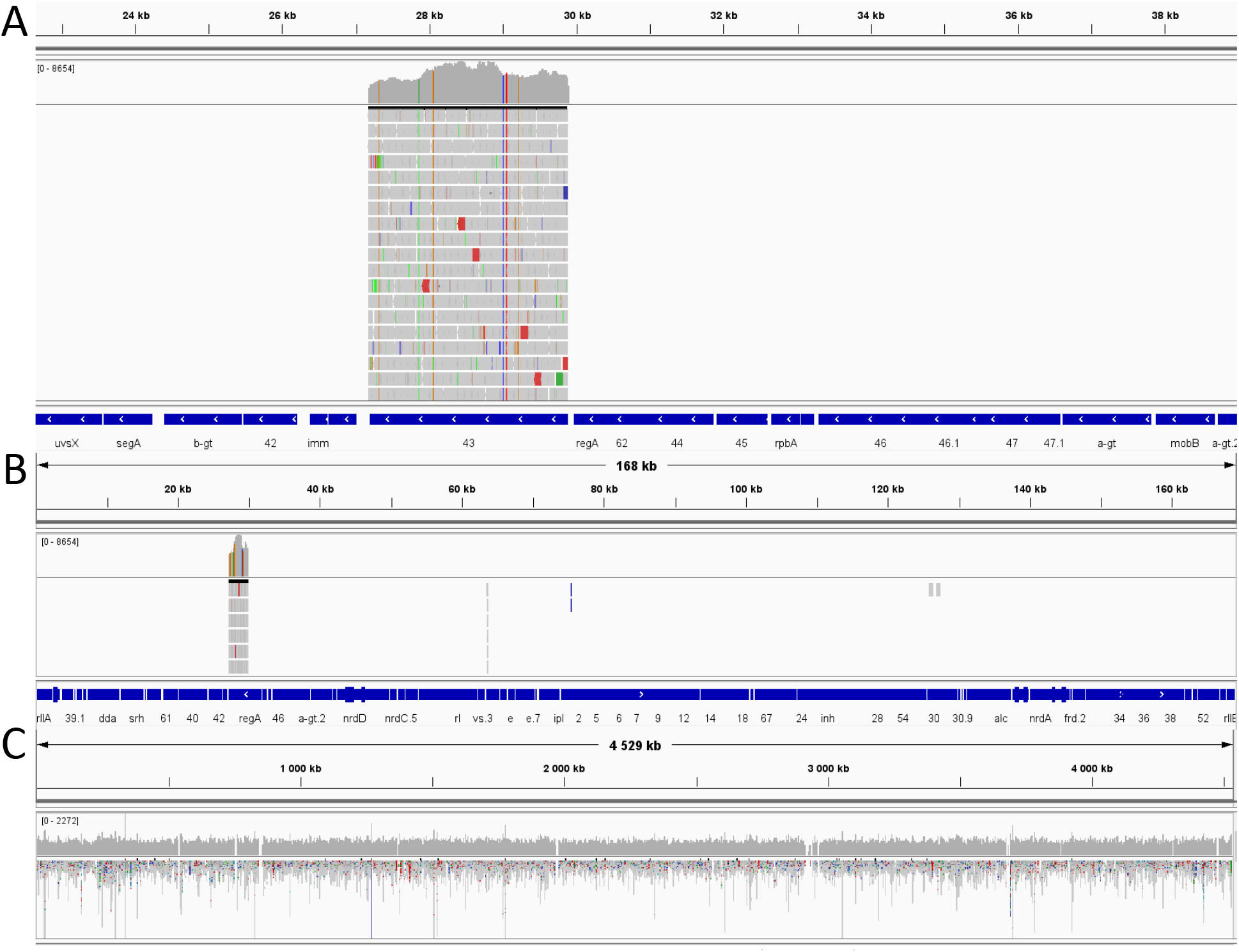
DIY library RNAseq data for the “15 ng” sample (Table 1) mapped to: *Enterobacteria* phage T4 genome (NCBI RefSeq Accession NC_000866.4), zoomed data and the whole-genome charts (respectively, **A** and **B**); *E. coli* K12 BL21 (DE3) genome, log scale (**C**). The sequencing data correspond to a single gene of T4 (“43”), with the rest of the T4 genome totally clean. The uniform coverage of *E. coli* genome implicates it as the source of contaminating DNA, which is all the more likely given its role as production host for the T4 DNA polymerase protein.

The coverage of T4 phage genome was strictly limited to a single gene “43” (Figure 2 A and B) encoding the DNA-directed DNA polymerase (NCBI RefSeq Accession NP_049662.1), a recombinant version of which we used in the ERAT step of the DIY library preparation protocol. The uniform coverage obtained for *E. coli* reference genome (random spikes do not represent any particular genes) implicates these bacteria, a production host organism for T4 DNA polymerase, as the source of the observed contamination (Figure 2 C). Thus, we identified the contamination source while demonstrating that the rest of T4 enzymes used in the protocol were DNA-free.

### Replacement of the contaminated enzyme

To solve the problem, we replaced T4 DNA polymerase in the ERAT step (see Methods) with *E. coli* DNA polymerase I large (Klenow) fragment. To test the eligibility, we set up a comparative experiment using 150 ng of total RNA extracted from HepG2 human cells for cDNA synthesis; the reaction was run in technical replicates designated 1 and 2. The double-stranded cDNA was purified on SPRI magnetic beads (see Methods) with a double-volume elution split into two equal volumes. These halved cDNA samples were further processed in parallel, with either T4 DNA polymerase or Klenow fragment used in the ERAT step (see Methods), all other steps being identical. Thus, the experiment involved direct comparison of RNAseq libraries prepared from 75 ng of total RNA with the use of either T4 DNA polymerase or Klenow fragment for ERAT. Table 3 summarizes the results of Kraken2 analysis and STAR mapping obtained with the two alternative enzymes. It shows that fractions of the data mapped to human genome, *E. coli* chromosome, and T4 DNA polymerase-encoding gene for technical replicates are identical. Importantly, contamination of the data with *E. coli* and T4 sequences is eliminated by the enzyme replacement. Most satisfactorily, 97% of NGS data obtained with “Klenow” libraries are mapped to human genome. To exclude the possible degenerating effect of the reduced exonuclease activity of Klenow fragment compared with T4 DNA polymerase, we evaluated the data complexity by counting duplicated reads and found no significant increase in the redundancy for “Klenow” libraries compared with “T4” libraries.

**Table 3.** Characteristics of RNAseq libraries made with use of T4 DNA polymerase and Klenow fragment. Comparison of RNAseq data for the libraries prepared using T4 DNA polymerase *vs*. Klenow fragment; 1 and 2 are two technical replicates of the experiment.

| Sample (technical repeat #) | 1 | 2 | 1 | 2 |
| --- | --- | --- | --- | --- |
| Enzyme used for ERAT | T4 pol | T4 pol | Klenow | Klenow |
| Kraken2 Homo sapiens, % | 78.2 | 79.2 | 93.7 | 94.2 |
| Kraken2 Enterobacteriaceae, % | 8.05 | 7.71 | 0.15 | 0.11 |
| Kraken2 Tequatrovirus, % | 5.6 | 5.2 | 0 | 0 |
| STAR mapped to human, % | 80.6 | 81.1 | 97.0 | 96.3 |
| Duplicates, % | 22.2 | 19.5 | 23.7 | 21.9 |

The 3% of non-useful data obtained with “Klenow” libraries is a minor fraction often present even in WGS datasets. According to Kraken2, the unmapped 3% come from different sources, with 0.7% roughly attributed to Proteobacteria and 0.32 % belonging to Proteus phage VB_PmiS-Isfahan (NCBI Taxonomy ID:1969841; [6]) whose host bacterium *P. mirabilis* inhabits soil and water. A significant proportion of non-useful data may represent sequencing artifacts derived from adapter/primer multimers, etc. [7].

## Discussion

We used a combination of commonly available tools and methods to identify key parameters of RNAseq data and perform sufficient data quality control. As a result, we identified contamination of sequencing libraries with foreign DNA of bacterial/viral origin, apparently related to impurities in certain ingredients of our DIY RNAseq library preparation protocol. The source was traced to T4 DNA polymerase contaminated with the host strain DNA. The immediate solution was to replace the contaminated T4 DNA polymerase with Klenow fragment (SibEnzyme E325). The results indicate that Klenow fragment can be used as a replacement for T4 DNA polymerase in DIY RNAseq protocols. Overall, the reported experience illustrates the utility of common approaches in overcoming obstacles while creating DIY protocols for NGS.

## Methods

### RNA extraction

Total RNA was isolated using either TRIzol™ Reagent (Invitrogen 15596018), RNeasy Mini Kit (Qiagen 74104), or AllPrep DNA/RNA Mini Kit (Qiagen 80204). DNAse treatment was performed with DNase I RNase-free (Thermo Scientific EN0525). Quality control by electrophoresis was performed for each sample, and only samples with clearly visible rRNA bands were taken for further processing. The lack of detectable DNA in the DNase-treated samples was confirmed by qPCR assay (QuantumDNA-Set, Evrogen QS004). RNA concentrations were measured using Qubit™ RNA HS Assay Kit (Invitrogen Q32852).

### Reverse transcription

Reverse transcription was performed using MMLV RT kit (Evrogen SK021). The synthesis was primed with oligo(dT). All reactions were performed for 1 hour at 42 °C and stopped by a 10-minute incubation at 70 °C. The cDNA samples were stored at −20 °C for no longer than 2 days before the next processing step.

### DNA fragmentation

As priming the reverse transcription with oligo(dT) implies preserved RNA integrity, the fragmentation was applied to cDNA. We used a Covaris S220 ultrasonicator according to the manufacturer’s recommended settings.

### Second strand cDNA synthesis

Second strand cDNA synthesis was performed using a mixture of DNA Polymerase I (Invitrogen 18010017) and RNAse H (Invitrogen 18021014) at 15 °C for 2.5 hours [8].

### Library preparation and sequencing

The libraries were prepared as described elsewhere [9] with slight modifications. The adapters, designed specifically for MGI sequencing, were diluted 10–20 fold from the original concentration to avoid formation of adapter-adapter ligation products, and the PCR cycle number was increased to 15–16. The end repair and A-tailing step (ERAT) was performed using the following enzymes: T4 polynucleotide kinase (SibEnzyme E312), T4 DNA polymerase as a default option or *E. coli* DNA polymerase I Klenow fragment (SibEnzyme E325), and Taq DNA polymerase (Evrogen PK113L). The ligation step involved T4 DNA ligase (SibEnzyme E320). The PCRs were carried out with KAPA HiFi PCR Kit (KAPABiosystems KR0368).

All DNA purification steps were performed with SPRI HighPrep™ PCR system (MagBio AC-60500) according to the manufacturer’s protocol using an appropriate beads-to-DNA volume ratio.

The sequencing was run on MGISEQ-2000 sequencer (MGI DNBSEQ-G400) in paired-end mode (× 100) according to the manufacturer’s protocols. At least 3 × 10^6^ reads were generated for each sample to perform QC analysis.

### Data processing

Data processing included the first step of FastQC [10] analysis, followed by downsampling of the read number to 3 × 10^6^ for each sample using Seqtk [11], elimination of imbalanced nucleotides, quality trimming and adapter clipping with Trimmomatic [12], finalized by another FastQC to confirm quality and nucleotide balance for the reads.

The mapping was performed on GRch38.p13 reference genome (NCBI RefSeq Assembly Accession GCF_000001405.39) using STAR aligner 2.7.9a [13]. All manipulations with .sam and .bam files were carried out in SAMtools software [14]. Picard [15] MarkDuplicates was used to count duplicates in .bam files. The duplicates were not removed from the data. The assignment to genomic features, including rRNA, was done in the featureCounts software [16]. For 18s, 28s and 5s rRNA, the counts were summed up and divided by the total number of reads. For the exome, introns and lncRNA counts, a .gtf was generated based on RefSeq .gtf files sorting all exons, introns, and lncRNAs into single eponymous features. The mitochondrial transcript (mtRNA) counts encompassed all reads mapping to mitochondrial genome. Non-transcribed (intergenic) regions coverage was estimated through a .gtf file containing no annotated genes or pseudogenes. Removal of pseudogenes allows to drop intergenic regions that are highly homologues to actually transcribed regions. Thus, it prevents overstatement for intergenic coverage.

Kraken2 software [4] was used to estimate species representation in samples that might contain not only RNA of the studied biological object (in our case *H. sapiens*), but also genomic/transcriptomic sequences of other species.

The T4 and *E. coli* genome data mapping was accomplished with BWA-MEM software [5]. The percentage of mapped sequences was estimated using SAMTools flagstat options. For T4 phage genome, coverage density was assessed visually in IGV [2]; regions with non-zero coverage were subject to exact characterization.

